# *Drosophila* clock cells use multiple mechanisms to transmit time-of-day signals in the brain

**DOI:** 10.1101/2020.10.26.353631

**Authors:** Annika F. Barber, Shi Yi Fong, Anna Kolesnik, Michael Fetchko, Amita Sehgal

## Abstract

Regulation of circadian behavior and physiology by the *Drosophila* brain clock requires communication from central clock neurons to downstream output regions, but the mechanism by which clock cells regulate downstream targets is not known. We show here that the *pars intercerebralis* (PI), previously identified as a target of the morning cells in the clock network, also receives input from evening cells. We determined that morning and evening clock neurons have time of day dependent connectivity to the PI, which is regulated by specific peptides as well as by fast neurotransmitters. Interestingly, PI cells that secrete the peptide DH44, and control rest:activity rhythms, are inhibited by clock inputs while insulin-producing cells are activated, indicating that the same clock cells can use different mechanisms to drive cycling in output neurons. Inputs of morning cells to the *DILP2*+ neurons are relevant for the circadian rhythm of feeding, reinforcing the role of the PI as a circadian relay that controls multiple behavioral outputs. Our findings provide mechanisms by which clock neurons signal to non-clock cells to drive rhythms of behavior.

**Significance Statement:** Despite our growing understanding of how the fly clock network maintains free-running rhythms of behavior and physiology, little is known about how information is communicated from the clock network to the rest of the brain to regulate behavior. We identify glutamate and acetylcholine as key neurotransmitters signaling from clock neurons to the *pars interecerebralis* (PI), a clock output region regulating circadian rhythms of sleep and metabolism. We report a novel link between *Drosophila* evening clock neurons and the PI, and find that the effect of clock neurons on output neuron physiology varies, suggesting that the same clock cells use multiple mechanisms simultaneously to drive cycling in output neurons.

## Introduction

Many physiological and behavioral processes across organisms exhibit daily rhythms controlled by an internal circadian system. The temporal organization conferred by circadian clocks allows anticipation of environmental changes and the coordination of biochemical and physiological processes within and across cells and tissues. Work in *Drosophila* and other organisms has elucidated the molecular basis of circadian pacemaking in the brain, but we still lack a molecular understanding of how time of day information is relayed from brain clock circuitry to downstream output regions that regulate circadian physiology and behavior.

The *Drosophila* clock network consists of ~150 neurons that express the core clock transcription-translation feedback loop genes(1). Of these, the ventrolateral neurons (LNvs) and dorsal lateral (LNd) clock neurons are important pacemakers for driving circadian rhythms in constant darkness, and coordinating the activity of other neurons in the clock network (2–5). Indeed, robust rhythms are an emergent property of the clock network as a whole (6–10). The rhythmic pattern of locomotor activity in light:dark (LD) cycles, which consists of morning and evening peaks that anticipate dawn and dusk respectively, is also attributed to specific clock cells. The LNvs act through the neuropeptide pigment-dispersing factor (PDF) to drive the morning anticipatory bout of locomotor activity while LNd clock neurons are termed evening neurons due to their role in regulating evening activity (3, 11, 12). LNd neurons are a molecularly heterogeneous population with subpopulations expressing different signaling molecules including the classical transmitter acetylcholine and the neuropeptides ITP, NPF and sNPF (13–16) DN1 dorsal neurons, which receive input from LNvs, also promote morning anticipation(17, 18), and exhibit highest activity in the pre-dawn and morning hours(12, 19). DN1 neurons express glutamate and the neuropeptide DH31, both of which have previously described roles in regulating circadian rhythms (16, 17, 20). The diverse array of signaling molecules expressed by neurons in the clock network offers many possibilities for signaling to brain regions that lack their own clocks but are important for the generation of behavioral rhythms (output cells).

A major clock output region in *Drosophila* is the *pars intercerebralis* (PI), a proto-hypothalamic region implicated in regulating circadian locomotor rhythms and peripheral cycling(21–26). Like the hypothalamus, the PI also controls aspects of sleep and feeding (27–31), and consists of different peptidergic cells. PI cells that express the neuropeptide diuretic hormone 44 (DH44), the fly ortholog of corticotropin-releasing factor, modulate rest:activity rhythms(22, 25). On the other hand, PI cells producing *Drosophila* insulin-like peptide 2 (DILP2) neurons are implicated in circadian gene expression in the fat body(23, 32), but have not been linked to behavioral rhythms.

As PI cells do not express the molecular clock machinery, time of day information must be relayed directly or indirectly from clock neurons to the PI. Previous work showed that the DN1 clock neurons project to two groups of PI neurons, the diuretic hormone 44 positive (*DH44*+) and the *Drosophila* insulin-like peptide 2 positive (*DILP2*+) groups. Hyperactivation of *DH44*+ neurons or RNAi knockdown of DH44 peptide is sufficient to ablate rest:activity rhythms in *Drosophila(22*), however silencing of DN1 neurons only weakens rest:activity rhythms(33). Thus, there must be additional upstream inputs to PI neurons that maintain rest:activity rhythms in the absence of DN1 signals. We hypothesized that robust circadian control of the PI would require inputs from both morning- and evening-active clock neurons, including perhaps long-distance signals directly from LNvs. We sought to identify additional clock groups signaling to the PI and also investigated the signaling molecules involved. We demonstrate that both *DH44*+ and *DILP2*+ neurons receive time-of-day dependent inputs from both CRY-negative LNds and DN1s. Surprisingly, clock neurons inhibit *DH44*+ neurons while activating *DILP2*+ neurons, indicating that the same clock cells use multiple mechanisms simultaneously to drive cycling in output neurons. We show also that morning clock cells and PI-secreted DILPs are required for rhythms of feeding.

## Results

### LNd and DN1 clock neurons have time of day dependent connectivity to the PI

Previous work has demonstrated that DN1 clock neurons have physical and functional connectivity to *DH44*+ and *DILP2*+ cells of the PI (22, 23), however it is unlikely that a single morning-active clock population is sufficient to confer robust circadian regulation of PI neuron activity. We used GFP reconstitution across synaptic partners (GRASP) to test the hypothesis that the LNd clock neurons are also presynaptic to PI neurons using the dvPDF-Gal4 driver with PDF-Gal80 (34–36). We tested connectivity via GRASP in brains dissected in the morning (ZT 0-4) and found evidence of physical connections from LNd neurons to both *DH44*+ and *DILP2*+ PI neurons (Fig. 1 A, B).

**Fig 1.**
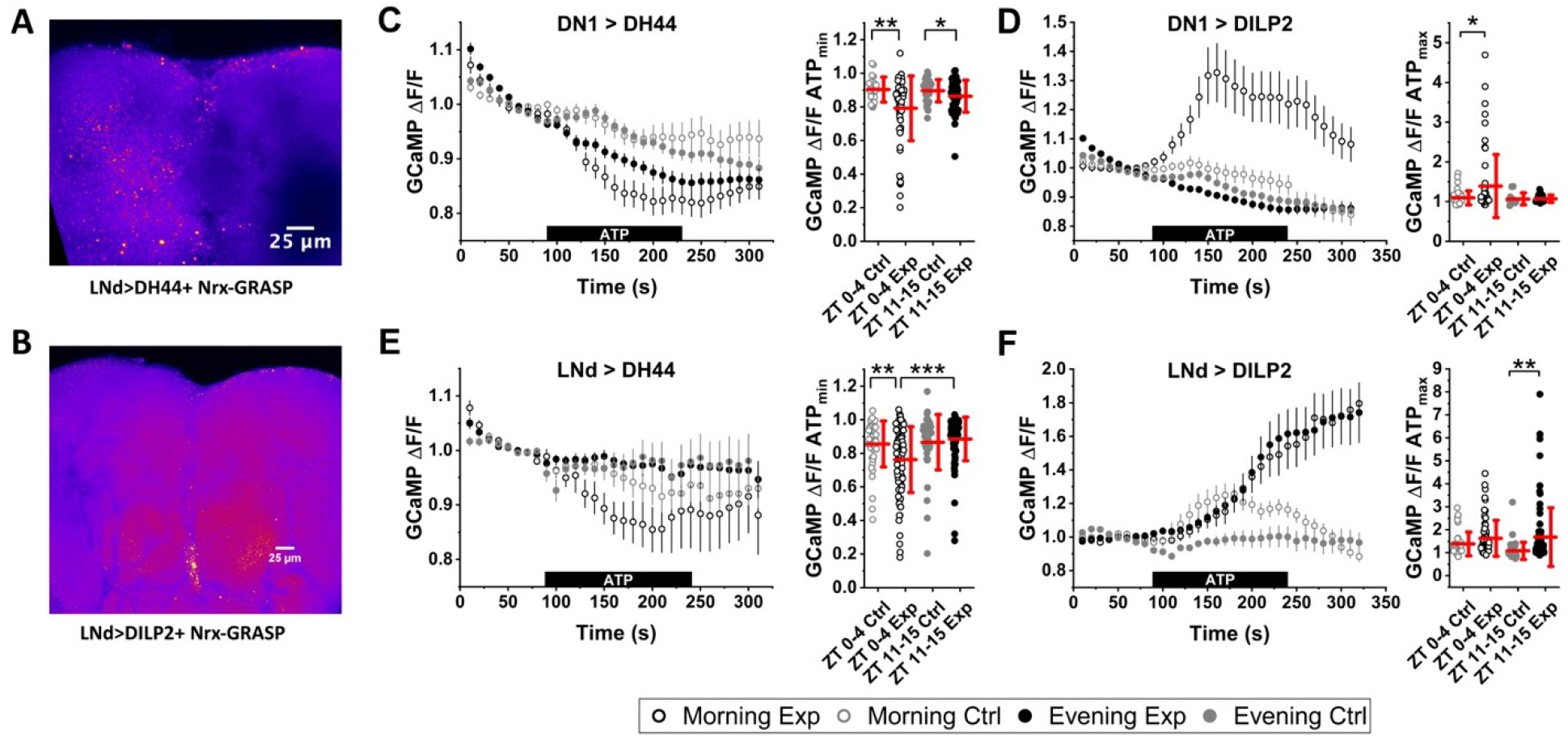
Clock neurons have time-of-day-dependent functional connectivity to the PI. (*A*) nrx-GRASP between CRY-negative LNd clock neurons and *DH44*+ PI neurons. (*B*) nrx-GRASP between LNd clock neurons and *DILP2*+ PI neurons. (*C) Left*: GCaMP6m signal over time in *DH44*+ neurons during activation of DN1 neurons. ZT 0: n = 66 cells, 15 brains, ZT 12: n = 42 cells, 11 brains. Black bar indicates timing of ATP application. Data are represented as mean ± SEM. *Right*: Minimum GCaMP change (ΔF/F), points represent values in individual cells, lines represent the mean ± SD for indicated genotypes and timepoints. ZT 0 control: n = 22 cells, 7 brains, ZT 12 control: n = 35 cells, 8 brains. (*D*) *Left*: GCaMP6m signal over time in *DILP2*+ neurons during activation of DN1 neurons. *Right*: Maximum GCaMP ΔF/F for each cell. Data plotted as described in (C). ZT 0: n = 60 cells, 12 brains, ZT 12: n = 45 cells, 10 brains., ZT 0 control: n = 39 cells, 7 brains, ZT 12 control: n = 23 cells, 5 brains. (*E*) *Left*: GCaMP6m signal over time in *DH44*+ neurons during activation of LNd neurons. *Right*: Minimum GCaMP ΔF/F for each cell. Data plotted as described in (C). ZT 0: n = 90 cells, 23 brains, ZT 12: n = 83 cells, 30 brains., ZT 0 control: n = 45 cells, 14 brains, ZT 12 control: n = 44 cells, 13 brains. (*F*) *Left*: Average GCaMP6m signal over time in *DILP2*+ neurons during activation of LNd neurons. *Right*: Maximum GCaMP ΔF/F for each cell. Data plotted as described in (C). ZT 0: n = 73 cells, 9 brains, ZT 12: n = 92 cells, 13 brains., ZT 0 control: n = 49 cells, 8 brains, ZT 12 control: n = 44 cells, 6 brains. * indicates p < 0.05, ** indicates p < 0.01 and *** indicates p < 0.001 by ANOVA followed by Tukey test for all panels.

To assay functional inputs from clock neurons to the PI, we used a calcium indicator-based stimulus-response assay (Fig. 1 C-F). In this assay, the ATP-gated cation channel P2X2 was expressed in clock neurons to allow specific depolarization by ATP application, while the genetically encoded calcium indicator GCaMP6m was expressed in PI neurons to allow monitoring of intracellular calcium levels before and after clock neuron stimulation in acutely dissected brains (37, 38). To control for effects of possible leaky P2X2 expression, or direct effects of ATP on calcium signaling in the PI, we expressed GCaMP6m in PI neurons of flies carrying *UAS-P2X2* but no GAL4 driver as a control.

Stimulation of clock neurons had divergent effects on *DH44*+ and *DILP2*+ neurons depending on time of day. In the morning (ZT 0-4), DN1 neuron activation resulted in a 21% ± 2% decrease in *DH44*+ cell GCaMP fluorescence, while in the evening (ZT 11-15) the 14% ± 1% reduction in the *DH44*+ GCaMP signal was indistinguishable from that in genetic controls (Fig. 1C). A time-dependent profile is also apparent for *DILP2*+ cells; DN1 neuron activation results in an 39% ± 10% increase in GCaMP fluorescence in the morning, and no change in the evening (Fig. 1D). It should be noted, however, that while the group response of both *DH44*+ and *DILP2*+ neurons to DN1 activation in the morning was significant, it was also highly heterogeneous, with some cells showing strong responses while others were indistinguishable from controls (Fig, 1C, D right panels). In contrast, the lack of response to DN1 activation in the evening in both cell groups was very consistent.

LNd neurons also show preferential morning modulation of *DH44*+ neurons (Fig. 1E), with 24% ± 2% inhibition of *DH44*+ GCaMP signaling in the morning and consistent lack of response in the evening. The *DILP2*+ neuron GCaMP response to LNd stimulation is heterogeneous in both the morning and the evening, with some neurons showing a large response to LNd activation at each time, while others show no response (Fig. 1F, right). Overall, in the morning there was a 58% ± 11% increase in *DILP2*+ cell GCaMP signal, however this was not significantly different from controls (43% ± 8%). In the evening there was a 68% ± 14% increase in DILP2 GCaMP signal, which was significantly different from controls (9% ± 6%). The increased GCaMP signal in morning controls may reflect the higher morning basal activity of *DILP2*+ neurons previously described(23). The heterogeneity of the *DILP2*+ neuron response to LNd stimulation obscures some aspects of the data when averaging. Thus, we used a 10% change in GCaMP6m signal during ATP application as a cutoff to delineate responding and non-responding neurons to allow us to compare the magnitude of response. The magnitude of the GCaMP ΔF/F increase in responding neurons was not significantly different between morning and evening time points at 87% ± 11% in the morning and 60% ± 9% in the evening (data not shown). The proportion of responding neurons varied only slightly by time of day, with 57% responding at ZT 0-4 and 65% responding at ZT 11-15.

### Application of clock neuron derived neuropeptides alters excitability of PI neurons

We next asked if neuropeptides produced by clock cells and implicated in various aspects of circadian rhythms signal to the PI to regulate behavioral rhythms. Diuretic hormone 31 (DH31), produced by dorsal clock neurons, regulates temperature preference rhythms through its canonical receptor, DH31-R, and also works in concert with PDF to regulate circadian locomotor rhythms through an unknown signaling pathway(39, 40). Bath application of 1 μM DH31 peptide onto acutely dissected GCaMP6m-expressing brains caused an acute increase in intracellular calcium in *DH44*+ neurons (19% ± 5%) (Fig. S1A, B). Also, 1μM DH31 had a heterogeneous effect on DILP2 neurons, with 22/52 neurons showing an increase in intracellular calcium as measured by GCaMP6m fluorescence (22% ± 3% c) and 30/52 neurons showing no change in GCaMP signal (2% ± 1 %) (Fig. S1 A, B). We previously reported that the *DILP2*+ neurons are heterogeneous in their response to signals from DN1s, with some neurons showing an increase in intracellular calcium and others showing no change (23).

Neuropeptide F (NPF), produced by LNd clock neurons, regulates free-running locomotor period and the amplitude of evening activity in a light-dark cycle, and also regulates circadian gene transcription in the fat body(14, 41). As with DH31, application of 1 μM NPF increased intracellular calcium in *DH44*+ neurons (12% ± 6%) (Fig. S1C, D). However, NPF had little effect on *DILP2*+ neurons (−6% ± 2%).

To examine roles for clock neuron derived neuropeptides in modulating circadian locomotor output through *DH44*+ neurons, we examined locomotor rhythms in DD after *DH44*-Gal4-driven RNAi knockdown of DH31 and NPF receptors. We saw no consistent change in total activity, period, or rhythm strength relative to parental controls in constant conditions; this is consistent with previous studies of DH31, which did not find effects on freerunning rhythms, and suggest that effects of NPF on behavior do not depend upon DH44 neurons (Fig. S2 A-C). Note that a behavioral difference would be considered significant only if the experimental line is significantly different from both parental controls. We also considered the possibility that PDF acts directly on the PI; although LNvs are not known to directly contact the PI, they project close to it in the protocerebrum and so limited diffusion of PDF is possible. As noted above, PDF signaling is implicated in freerunning rhythms and also in the maintenance of a characteristic morning-evening pattern in light:dark cycles; *pdfr* mutants have reduced morning anticipation and phase shifted evening anticipation in LD conditions (42). While knockdown of PDFR in DH44 cells did not affect rhythms in constant conditions, it reduced the evening peak of activity in a 12:12 light-dark cycle, (Fig. S3).

### Acetylcholine and glutamate are required for clock-to-PI signaling

Two major fast neurotransmitters are produced by the clock neurons upstream of the PI: glutamate from DN1 neurons, and acetylcholine from a subset of LNd neurons (13, 16, 20, 33). To examine the role of clock-secreted fast transmitters in the clock-to-PI circuit, we conducted the stimulus-response paradigm described above in the presence of specific neurotransmitter receptor blockers. Specifically, we asked if pre-treatment with a blocker attenuated the effect of P2X2 stimulation of upstream clock neurons on downstream PI intracellular calcium levels. All experiments were conducted in the morning, as this was the time of maximal PI neuron response for three out of four connections. We hypothesized that NMDAR or mGluR blockers would be most effective in attenuating the response of PI neurons to DN1 stimulation, while mAChR or nAChR blockers would attenuate the response to LNd stimulation. To our surprise, the most effective blocker was unique to each of the four connections tested, and did not necessarily correlate with the neurotransmitter believed to be released by the upstream clock population (Fig. 2).

**Fig. 2.**
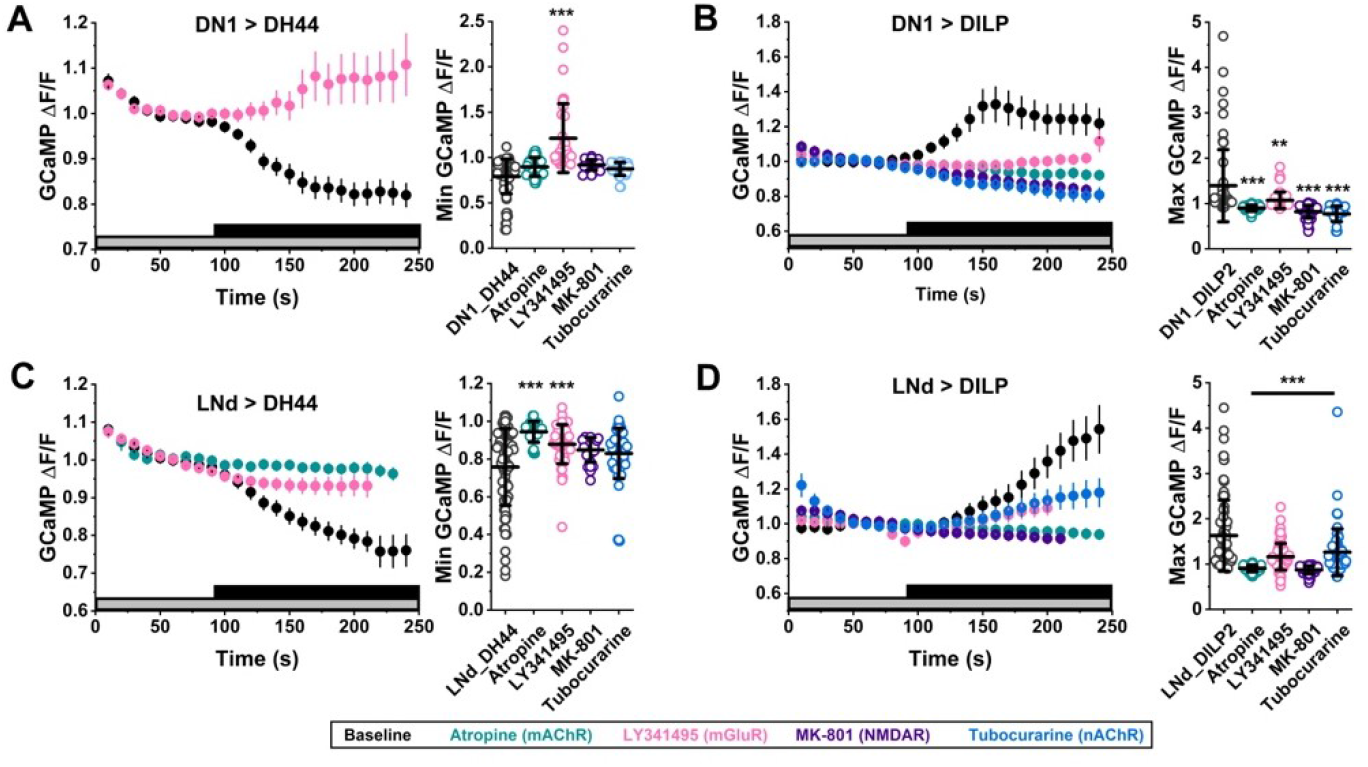
Acetylcholine and glutamate receptor blockers attenuate clock-to-PI signaling. (*A) Left*: GCaMP6m signal over time in *DH44*+ neurons during activation of DN1 neurons in the presence of the mGluR blocker LY341495. Control data for ATP activation only are replotted from Fig. 1A (black). Black bar indicates timing of ATP application; grey bar indicates timing of blocker application. Data are represented as mean ± SEM. *Right*: Minimum GCaMP change (ΔF/F), points represent values in individual cells, lines represent the mean ± SD for indicated receptor blocker. Control data for ATP activation only are replotted from Fig. 1C (black). N_atropine_ = 28 cells from 8 brains. N_LY341495_ = 38 cells from 9 brains. N_MK-801_ = 21 cells from 6 brains. N_tubocurarine_ = 31 cells from 8 brains. (*B) Left*: Data plotted as in (A) for *DILP*+ neurons during activation of DN1 neurons in the presence of the mGluR and AChR blockers. Control data for ATP activation only are replotted from Fig. 1D (black). *Right*: Maximum GCaMP change (ΔF/F) for individual cells. N_atropine_ = 40 cells from 8 brains. N_LY341495_ = 50 cells from 8 brains. N_MK-801_ = 51 cells from 9 brains. N_tubocurarine_ = 22 cells from 4 brains. (*C) Left*: Data plotted as in (A) for *DH44*+ neurons during activation of LNd neurons in the presence of the mGluR and AChR blockers. Control data for ATP activation only are replotted from Fig. 1E (black). *Right*: Minimum GCaMP change (ΔF/F) for individual cells. N_atropine_ = 19 cells from 10 brains. N_LY341495_ = 48 cells from 8 brains. N_MK-801_ = 48 cells from 16 brains. N_tubocurarine_ = 46 cells from 13 brains. (*D) Left*: Data plotted as in (A) for *DILP*+ neurons during activation of LNd neurons in the presence of the mGluR and AChR blockers. Control data for ATP activation only are replotted from Fig. 1F (black). *Right*: Maximum GCaMP change (ΔF/F) for individual cells. N_atropine_ = 52 cells from 8 brains. N_LY341495_ = 62 cells from 9 brains. N_MK-801_ = 72 cells from 10 brains. N_tubocurarine_ = 62 cells from 10 brains. All data were collected at ZT 0-4. Asterisks indicate comparisons between each drug treatment and no-drug control (far left); ** indicates p < 0.01 and *** indicates p < 0.001 by ANOVA followed by Tukey test for all panels.

For the DN1-to-DH44 connection, all blockers tended to attenuate the reduction in intracellular calcium caused by DN1 neuron stimulation. However, only the mGluR blocker LY341495 caused a statistically significant change in the GCaMP response (Fig 2A). In some instances, LY341495 uniquely reversed the effect of DN1 neuron stimulation, such that we observed an average increase in *DH44*+ neuron intracellular calcium in the presence of the blocker (Fig. 2A).

For the DN1-to-DILP2 connection, all blockers tested significantly attenuated the increase in intracellular calcium caused by DN1 neuron stimulation. The nAChR blocker (tubocurarine), mAChR blocker (atropine) and NMDAR blocker (dizocilpine) all completely attenuated the DH44 GCaMP response to DN1 neuron stimulation (Fig 2B). Administration of the mGluR blocker LY341495 substantially reduced the DILP2 GCaMP response, but was slightly less effective than the other blockers (Fig. 2B).

For the LNd-to-DH44 connection, all blockers tested trended toward an attenuation in the reduction in intracellular calcium caused by LNd neuron stimulation. However, only the mAChR blocker (atropine) and mGluR blocker (LY341495) resulted in a statistically significant attenuation of the DH44 GCaMP response to LNd simulation (Fig. 2C).

For the LNd-to-DILP2 connection, all blockers tested significantly attenuated the increase in intracellular calcium caused by LNd neuron stimulation. Under baseline conditions, the *DILP2*+ neuron response to DN1 stimulation by ATP was heterogeneous, but resulted in an average increase in GCaMP fluorescence (Fig. 1D) In the presence of mAChR and NMDAR blockers (atropine and MK-801) the ATP-induced increase in intracellular calcium was completely blocked (Fig. 2D). The mGluR and nAChR blockers (LY341495 and tubocurarine) significantly reduced the ATP-induced increase in intracellular calcium compared to controls, but cells still responded with an increase in calcium above baseline (Fig. 2D).

To examine roles for clock neuron derived glutamate or acetylcholine modulation of circadian output through *DH44*+ neurons, we examined locomotor rhythms in DD after *DH44*-Gal4-driven RNAi knockdown of acetylcholine and glutamate receptors (Fig. S4). mAChR-A knockdown with two RNAi lines resulted in a significant increase in period, primarily due to outlier effects (Fig. S4B). One mGluR RNAi knockdown line lengthened period and reduced rhythmicity, but this was not recapitulated by other RNAi lines against the same target (Fig. S4C). In general, we conclude that knockdown of any single glutamate or acetylcholine receptor in DH44 neurons is not sufficient to induce alteration in behavioral rhythms, though that does not rule out participation of these receptors in concert with other redundant signaling mechanisms.

### Clock signaling through DILP2 neurons alters feeding rhythms and food intake

Our previous work further demonstrated that *DILP2*+ neurons exhibit diurnal rhythms of electrical activity in ad lib feeding conditions, and that altering the timing of feeding alters *DILP2*+ neuron activity(23). We hypothesized that despite the heretofore ambiguous role of insulin in feeding in flies, that insulin may be a circadian output molecule regulating rhythmic feeding. We used the activity recording CAFE (ARC) assay (43) to quantify single-fly feeding for 72 hours, with the first day in a 12:12 LD cycle and the following two days in constant darkness (DD). We first validated that our wild-type iso31 flies show rhythmic feeding and that this rhythm is lost in the absence of a functional clock due to the loss of the *period* gene (*per*^01^) (Fig. 3A, B). Wild-type flies exhibit feeding rhythms in LD that are maintained in DD. *per*^01^ flies lack feeding rhythms in both LD and DD, despite the fact that *per*^01^ flies exhibit rhythmic locomotor activity in LD(44). Additionally, *per*^01^ flies ate significantly more food per day than WT flies throughout the assay (Fig. 3B).

**Fig. 3.**
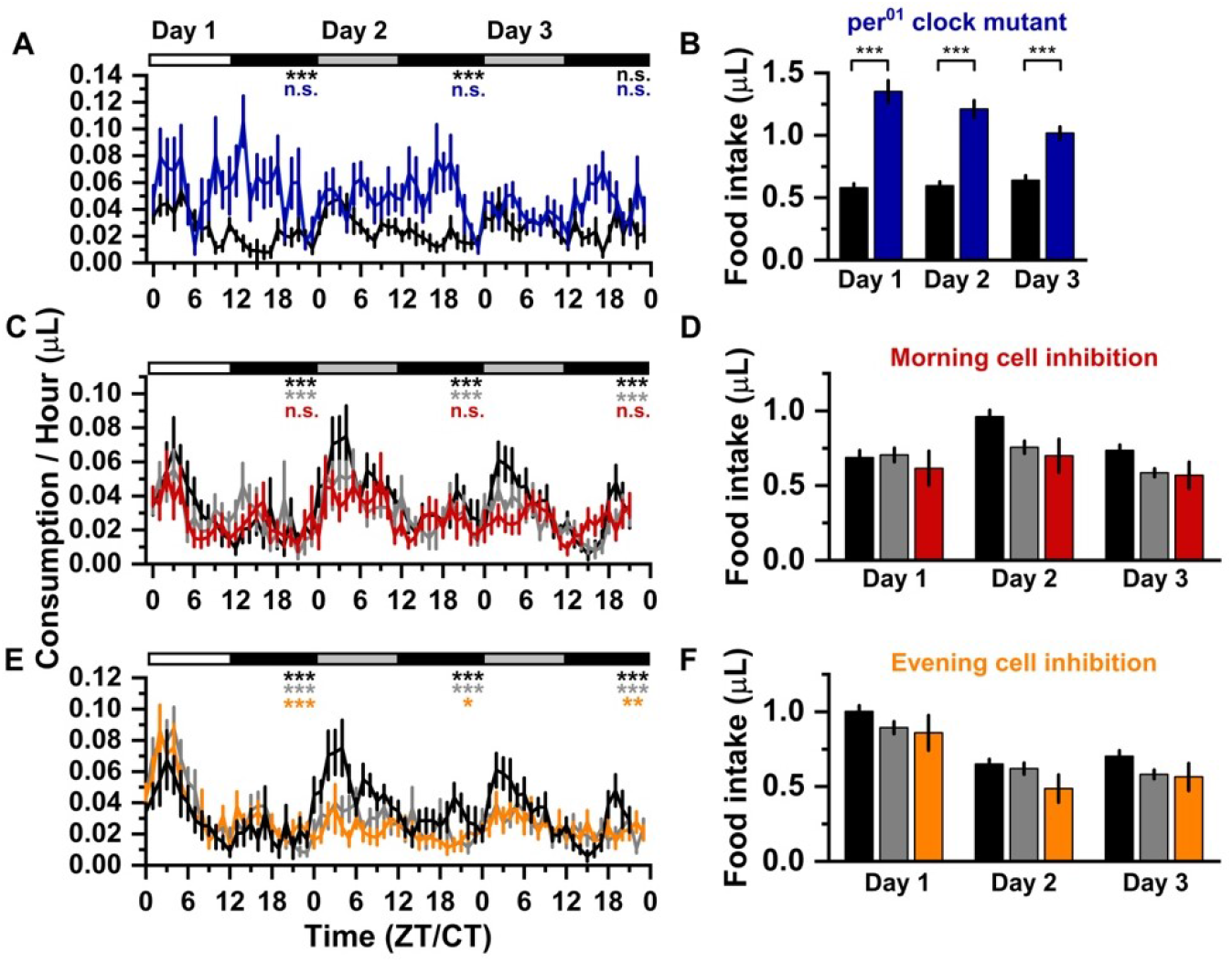
The clock controls daily feeding rhythms via *PDF*+ morning cells. (*A*) Average consumption in μL per fly per hour for wild-type (black, N = 23) and *Per*^01^ clock mutant (blue, N = 20) over three days. Error bars indicate SEM. Greyscale bars at the top indicate light conditions. White: lights-on, light grey: lights off (subjective day), dark grey: lights-off (night). JTK_cycle p-values were calculated by 24-hour day for each genotype, significance indicated below lighting bars. P_JTK_<0.001 = ***; P_JTK_>0.05 = arr. (arrhythmic). (*B*) 12-hour food intake in μL during the day (ZT/CT 0-11) vs. night (ZT/CT 12-23) for flies shown in (A). Day vs. night feeding was compared within each genotype, and 24-hour food intake was compared between genotypes by Student’s t-test. * indicates p < 0.05, ** indicates p < 0.01 and *** indicates p < 0.001. (*C*) Average consumption per fly per hour as in (A) for PDF-Gal4 > UAS-Kir2.1 (red, N = 24), Gal4 control (black, N = 25) and UAS control (grey, N = 25). (*D*) 12-hour food intake for flies shown in (C), plotted as in (B). There were no significant differences in 24-hour food intake between genotypes. (*E*) Average consumption per fly per hour for as in (A) for dvPDF-Gal4; PDF-Gal80 > Kir2.1 (mustard, N = 31), Gal4 control (black, N = 31) and UAS control (grey, N = 31). (*F*) 12-hour food intake for flies show in (E), plotted as in (B). There were no significant differences in 24-hour food intake between genotypes.

To determine the role of clock populations in regulating feeding rhythms, we examined the contribution of *pdf*+ “morning” cells and LNd “evening” cells to rhythmic food intake (Fig. 3 C-F). Suppression of morning cell activity by expression of the hyperpolarizing inwardly rectifying potassium channel Kir2.1 resulted in a loss of feeding rhythms in both LD and DD conditions, while controls remained rhythmic in both conditions (Fig. 3C). Total daily food intake was not altered by *pdf*-Gal4-driven expression of Kir2.1 (Fig. 3D). Suppression of evening cell activity, however, did not result in a loss of feeding rhythm in LD or DD and did not change daily food intake compared to controls (Figs. 3 E, F).

To assess the role of insulin-like peptides in regulating circadian feeding, we examined DILP2 and DILP2,3 insertion mutants (ΔDILP2 and ΔDILP2,3) in the ARC assay (Fig. 4). In the DILP2 mutant (Fig. 4A, B), rhythmic feeding was maintained in constant darkness, though the timing of the daily feeding peak was delayed relative to controls, from CT3 to CT6. There was no significant different in 24-hour food intake between ΔDILP2 mutant and controls. Because loss of one or more ILPs can result in compensatory upregulation of the remaining ILPs (45), the contribution of PI-derived ILPs is best-studied using a triple knockout of ILPs 2, 3 and 5. However, the ΔDILP2,3, 5 flies have multiple metabolic and reproductive defects that make them unsuitable for use in the feeding assay, hence we used ΔDILP2,3 flies. Loss of both DILPs 2 and 3 resulted in not only a loss of rhythm, but also a substantial reduction in total food intake (Fig. 4C, D). Control flies ate 0.82 ± 0.06 μL/day, while ΔDILP2,3 flies ate 0.51 ± 0.03 μL/day. While ΔDILP2,3 flies maintain a low amplitude rhythm in LD, this rhythm in immediately dampened in DD and is not detectable by JTK_cycle.

**Fig. 4.**
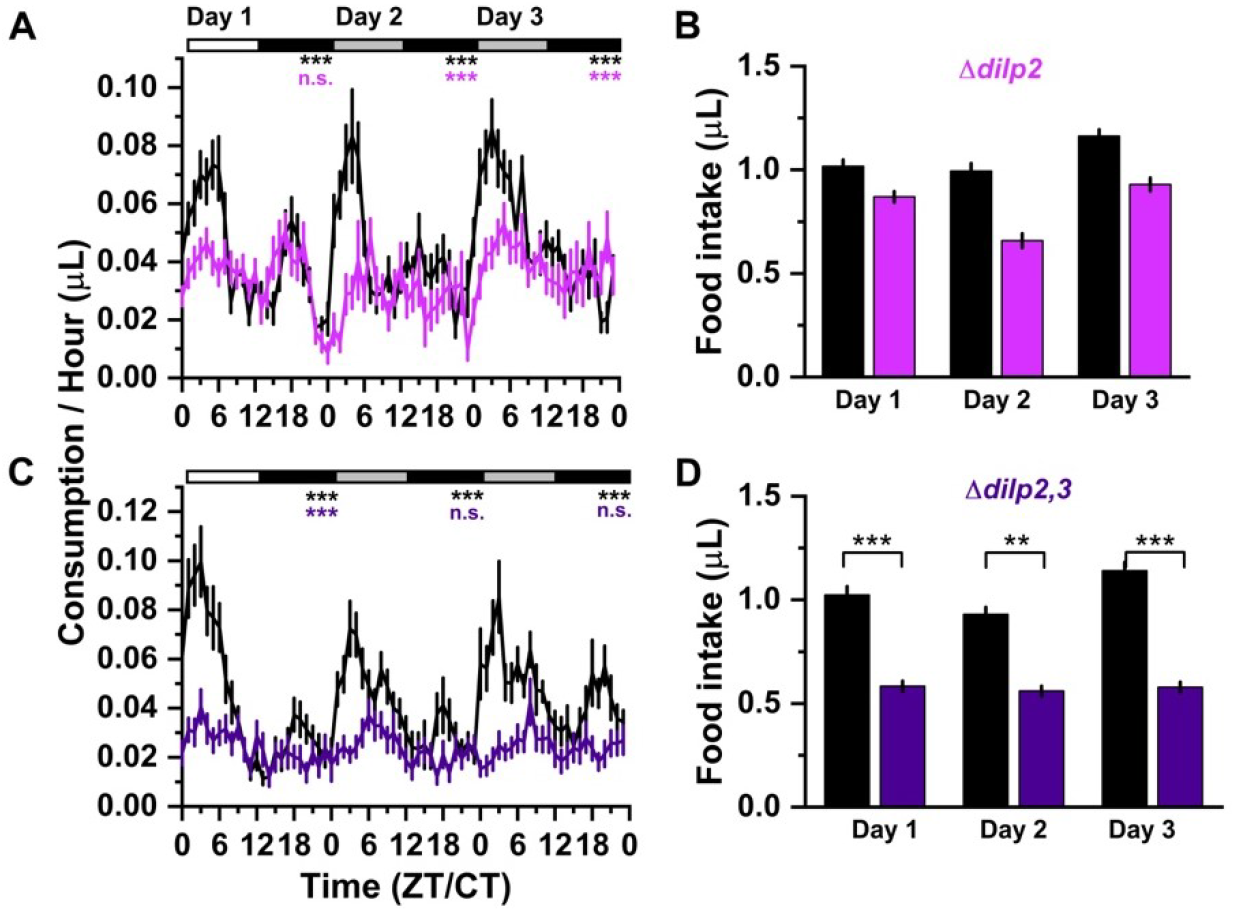
Brain-derived insulin regulates timing and amount of food intake. (*A*) Average consumption in μL per fly per hour for *DILP2*^-/+^ control (black, N = 54) and *DILP2*^-/-^ insertion mutant (magenta, N = 69) over three days. Error bars indicate SEM. Greyscale bars at the top indicate light conditions. White: lights-on, light grey: lights off (subjective day), dark grey: lights-off (night). JTK_cycle p-values were calculated by 24-hour day for each genotype, significance indicated below lighting bars. P_JTK_<0.001 = ***; P_JTK_>0.05 = arr. (arrhythmic). (*B*) 12-hour food intake in μL during the day (ZT/CT 0-11) vs. night (ZT/CT 12-23) for flies shown in (A). Day vs. night feeding was compared within each genotype, and 24-hour food intake was compared between genotypes by Student's t-test. * indicates p < 0.05, ** indicates p < 0.01 and *** indicates p < 0.001. There were no significant differences in 24-hour food intake between genotypes. (*C*) Average consumption per fly per hour for as in (A) for *DILP2,3*^-/+^ control (black, N = 67) and *DILP2,3*^-/-^ insertion mutant (purple, N = 66). (D) 12-hour food intake for flies show in (C), plotted as in (B), with statistical comparisons for 24-hour consumption above.

**Fig. 5.**
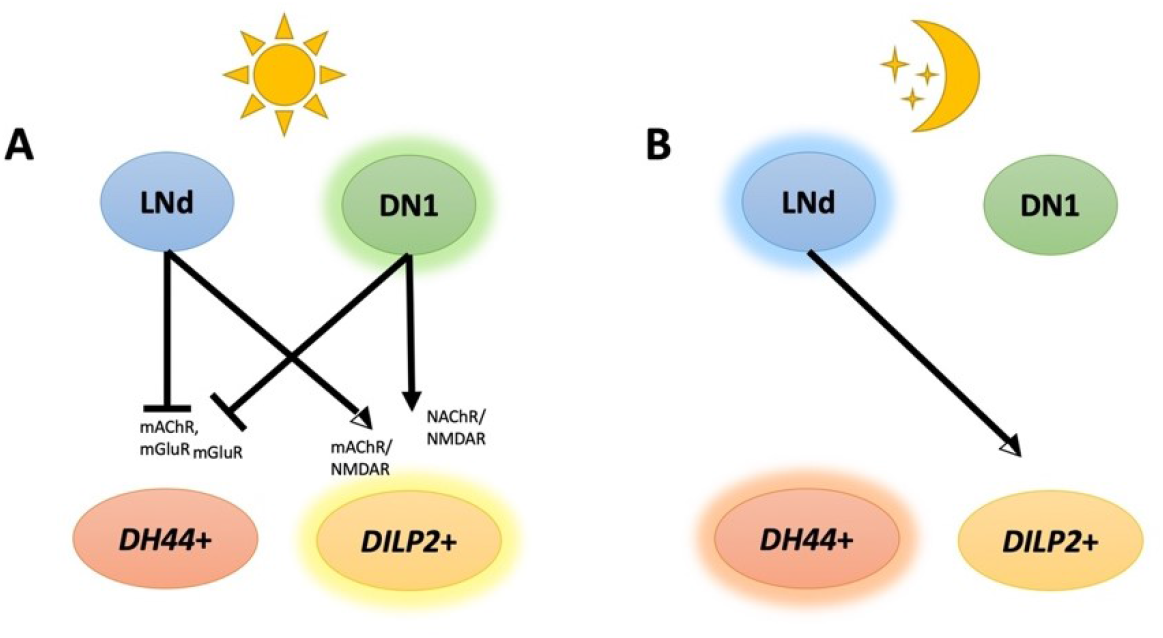
Model for time-of-day dependent modulation of PI neurons by clock populations. (*A*) In the morning, LNd and DN1 clock provide inhibitory signals to *DH44*+ neurons and excitatory signals to *DILP2*+ neurons. (B) At night, LNd neurons continue to provide excitatory signals to *DILP2*+ neurons, but no longer inhibit *DH44*+ neurons. DN1 neurons provide no modulatory signal to either PI population at night.

## Discussion

Previous work demonstrated that the Drosophila PI is a proto-hypothalamic region that serves as a circadian output hub(22, 23, 25, 31); while inputs from DN1 clock neurons have been shown, the signaling molecules involved in transducing time-of-day cues from the clock to the PI have been described. Our data suggest that time-of-day information arrives at the PI from multiple clock neuron populations through cholinergic and glutamatergic signaling. Further, we identify a new output role for the *DILP2*+ neurons of the PI in regulating circadian feeding, which depends on inputs from morning active, but not evening active, clock neurons.

The response of PI neurons to clock neuron stimulation is time-of-day dependent. *DH44*+ neurons are inhibited – i.e. exhibit a reduction in intracellular calcium - by both DN1 and LNd neuron stimulation in the morning, with little to no effect of stimulation of clock neurons in the evening. Though LNds drive the evening peak of locomotor activity(8), LNd neuron stimulation is only capable of inhibiting *DH44*+ neurons in the morning. *DH44*+ neurons are evening active in both LD and DD conditions(24, 26), and while they regulate both the morning and evening activity peaks in LD, their contribution to the evening peak is larger(22, 25). It is likely that *DH44*+ neurons, which are implicated in several processes (22, 25, 27, 29), receive non-clock inputs that drive locomotor activity, and they require daytime silencing by the clock to inhibit locomotion outside the morning and evening activity peaks. Thus, silencing by DN1s may serve to delineate the morning peak of activity; on the other hand, early day silencing of DH44 neurons by LNds may be one mechanism by which LNds drive evening activity.

The response of *DILP2*+ neurons to DN1 stimulation is time-of-day dependent, while the response to LNd stimulation is not. The *DILP2*+ population is larger than the *DH44*+ population and is known to be heterogeneous in terms of basal activity and signaling molecule expression (23, 46, 47). While the functional relevance of LNd signaling to *DILP2*+ cells is unclear, the DN1 inputs likely contribute to the circadian rhythm of feeding. This is consistent with the higher firing of *DILP*+ cells in the morning and the morning peak of the feeding rhythm in w^1118^ flies(23, 48) as well as with our finding here that DILPs are required for rhythmic feeding. In further support of this idea, we report that feeding rhythms depend upon the activity of morning active LNvs, which signal through DN1s, but not evening-active LNds. Our finding of loss of feeding rhythms on yeast/sucrose diet with genetic ablation differs from the findings of Dreyer et al. (31) who observed maintenance of rhythmic feeding on sucrose-only food upon adult silencing of *DILP2*+ neurons. We speculate that developmental changes caused by genetic ablation of insulin(s) may result in disorganized feeding patterns in constant conditions, and/or that rhythmic protein feeding, but not sugar feeding, is regulated by brain insulins.

The finding that nAChR and mGluR signaling contribute to inhibition in one case (input from DN1 and LNds on to DH44 neurons) and activation in another (input from DN1 and LNds on to DILP cells) raises the question of what determines the nature of the effect. The most straightforward explanation would be a difference in the receptor subtypes expressed in the two cell groups. Alternatively, other signaling components specific to each cell type could account for the differential response. Regardless, it is clear that the actions of known neurotransmitters can be regulated by the clock in different ways to effect rhythmic output. As noted above, acetylcholine is secreted by the LNds and likely also DN1s, but to date, the only known source of glutamate in the clock network are the DN1s. This suggests that glutamatergic input from the DN1s can modulate the effect of LNds on PI neurons.

Despite the effect of clock cell-secreted neuropeptides on PI cells and the role of fast neurotransmitters in signaling from clock cells to the PI, knockdown of neuropeptide/neurotransmitter receptors in the PI has little effect on behavioral rhythms (Fig. S2 and data not shown). There are many possible explanations for lack of a phenotype—low mRNA levels remaining after knockdown (we verified knockdown) are sufficient for function, receptor subtypes act redundantly in the regulation of rhythms, knockdown in the PI alone is insufficient to yield a phenotype or these molecules affect physiological rhythms other than those of rest:activity. While DH31 and NPF have been implicated in the control of rest:activity rhythms, loss of acetylcholine and glutamate have not been linked to behavioral rhythms. Although knockdown of receptors for these molecules in the PI did not yield a phenotype, we saw reduced evening activity with DH44-driven knockdown of PDFR. This is in contrast to the effect of loss-of-functions *pdf* mutants in which evening activity remains high, but is shifted ~1 hour early(42). It is possible that DH44 cell-mediated reduced evening activity is compensated in *pdfr* mutants. PDF is secreted by LNvs, which are not known to synapse onto the PI, but do project to the dorsal brain, suggesting diffusion of PDF across a small region. Limited diffusion of PDF is supported by a previous study showing that over-expression of PDF in cells that project to the dorsal brain produces behavioral phenotypes(49).

These findings elucidate some of the complexity underlying circadian control in neural circuits. Through the use of neuropeptides and fast neurotransmitters coupled with time-of-day specific actions on downstream neurons, clock cells are able to drive multiple outputs. Proximity of output neurons controlling locomotor activity (*DH44*+) with those that control feeding (*DILP*+) likely allows for integration of different behaviors and contributes to organismal fitness.

## Materials and Methods

Detailed materials and methods are provided in SI materials and methods.

### Fly stocks

UAS/Gal4 lines and mutants used for behavior and immunohistochemistry are described in the SI appendix. See Table S1 for a list of the complete genotype for the animals used in each experiment.

### P2X2 Activation and GCaMP imaging

GCaMP6m imaging in response to P2X2 activation in acutely dissected brains was performed as previously described (23, 25). Fly entrainment procedures and detailed methods are described in the SI appendix.

### Drosophila activity monitoring assay

Rest:activity rhythm assays were performed with the Drosophila Activity Monitoring System (Trikinetics, Waltham MA) as described previously (50). Fly entrainment procedures and detailed methods are described in the SI appendix.

### Activity recording CAFE assay

Single-fly circadian feeding was assessed using the Automated Recording CAFE assay (43). Detailed methods are described in the SI appendix.

### Immunohistochemistry, GRASP and confocal microscopy

Fly lines, entrainment conditions, antibodies and detailed methods are described in the SI appendix.

### Statistical analysis

Sample sizes and statistical details of each experiment can be found in the figure legends. All statistical tests were performed in OriginPro 2020. Additional detailed statistical methods are provided in the SI appendix.

## Acknowledgements

We thank Jin-Hong Scarlet Park and Drs. William Ja, Robert Huber and Keith Murphy for their patient technical advice and assistance with the ARC assay. Thanks also to Dania Malik for support with creating R scripts for ARC data analysis and visualization. Stocks from the Bloomington Drosophila Stock Center (NIH P40OD018537) and Vienna Drosophila Resource Center were used in this study. This work was supported by the Howard Hughes Medical Institute and NIH R37 NS048471 (to A.S.) and NIH-NINDS K99/R00 NS105942 (to A.F.B.).

## Supplemental Information

### SI Materials and Methods

#### Fly Stocks and Maintenance

Flies were maintained on cornmeal-molasses medium at 25°C. The *w1118* iso31 strain was used as wild-type(51). When tested as controls, UAS and GAL4 fly lines were tested as heterozygotes after crossing to iso31. RNAi lines used in behavioral screens were from the Transgenic RNAi Project (TRiP) collection from the Bloomington *Drosophila* Stock Center (BDSC) and the P-element (GD) and phiC31 integrase (KK) RNAi collections from the Vienna *Drosophila* Resource Center (VDRC). See Table S1 for a list of the complete genotype for the animals used in each experiment.

#### P2X2 Activation and GCaMP imaging

Adult male and female flies 5-10 d old were entrained at least 3 days in a 12 h light: 12 h dark (12:12 LD) cycle prior to functional imaging experiments. Flies were anesthetized on ice and dissected in hemolymph-like saline (HL3) consisting of (in mM): 70 NaCl, 5 KCl, 1.5 CaCl2, 20 MgCl2, 10 NaHCO3, 5 trehalose, 115 sucrose, 5 HEPES, pH 7.1 (37). Imaging experiments were performed using a naked brain preparation in a small bath of HL3 in a perfusion chamber (AutoMate Scientific, Berkeley CA). The brain was stabilized under nylon fibers attached to a platinum wire frame. Solutions were perfused over the brain at a rate of ~5 mL/min with a gravity-fed ValveLink perfusion system (AutoMate Scientific). After 1.5 min of baseline GCaMP6m imaging, ATP was delivered to the chamber by switching perfusion flow from the channel containing HL3 to another channel containing 2.5 mM ATP in HL3, pH 7.1. ATP was perfused for 2.5 min, followed by a 1-min washout with HL3. In experiments with neurotransmitter receptor blockers, the blocker was dissolved in HL3 at the indicated concentration and delivered during the 1.5-min baseline; the same concentration of blocker was then applied with the 2.5 mM ATP for 2.5 min. Blockers used were: atropine (7.3 μm), LY341495 (35 μm), MK-801 maleate (7.5 mM), tubocurarine (20 μm). GCaMP6 calcium imaging was performed on a Leica TCS SP5 confocal microscope. Twelve-bit images were acquired with a 40 x /0.8 water immersion objective at 256 x 256 pixel resolution. Z-stacks were acquired every 10 s.

Image processing and measurement of fluorescence intensity was performed in FIJI as previously described (25). A sum-intensity Z-projection of each time step was used for analysis, and the StackReg FIJI plugin was used to correct for small x-y movements over time in the sum-projected image. Regions of interest (ROIs) were manually drawn to encompass individual GCaMP-positive cell bodies, and mean fluorescence intensities were measured from each ROI at each time point. For each cell, fluorescence traces over time were normalized using this equation: ΔF/F = (F_n_-F_0_)/F_0_, where F_n_ is the fluorescence intensity recorded at time point n, and F_0_ is the average fluorescence value during the 30-s baseline preceding ATP application. Maximum ΔF/F was calculated by subtracting the average ΔF/F in the 30 s preceding ATP delivery from the largest ΔF/F value during ATP application. Brains with cells that have unstable baselines were discarded from quantification. We used Student’s t-test for two-group comparisons and one-way analysis of variance (ANOVA) followed by Tukey’s *post hoc* analysis to compare differences in maximum and minimum ΔF/F between groups.

#### Immunohistochemistry, GRASP and microscopy

Brains from 5-10 d old male flies entrained at least 3 d in a 12:12 LD cycle were dissected in phosphate-buffered saline with 0.1% Triton-X (PBST). Brains were fixed in 4% formaldehyde for 20 min at room temperature. Brains were rinsed 3 x 10 min with PBST, blocked for 60 min in 5% normal donkey serum in PBST (NDST), and incubated in primary antibody diluted in NDST for > 16 hr at 4°C. Brains were rinsed 3 x 10 min in PBST, incubated 2 hr in secondary antibody diluted in NDST, rinsed 3 x 10 min in PBST, and mounted in Vectashield (Vector Laboratories). Primary antibodies used are rabbit anti-GFP at 2mg/mL (Thermo Fisher Scientific A-11122), rat anti-RFP at 1mg/mL (ChromoTek 5F8), and mouse anti-bruchpilot at 1:100 (Developmental Studies Hybridoma Bank nc82). Secondary antibodies used are FITC donkey anti-rabbit (Jackson ImmunoResearch 711-095-152), Cy3 donkey anti-rat (712-165-153), and Cy5 donkey anti-mouse (715-175-151) at 1:500. Eight-bit images were acquired using a Leica TCS SP5 laser scanning confocal microscope with a 40x/1.3 NA or 20x/0.7 NA objective and a 1-mm z-step size. Image processing and measurement of fluorescence intensity was performed in FIJI(52). A maximum intensity Z-projection of each brain was used for analysis. Experimental GRASP intensity was compared to negative controls. We scored a GRASP signal as positive when more than half of brains within a time-point presented a reconstituted GFP signal.

#### Behavior experiment: Circadian rest:activity rhythms

Rest:activity rhythm assays were performed with the Drosophila Activity Monitoring System (Trikinetics, Waltham MA) as described previously (50). Flies were entrained to a 12:12 LD cycle for > 3 days at 25° C. 5 d old male flies were individually placed into glass tubes with 5% sucrose / 2% agar food and monitored in constant darkness (DD) for 7 d at 25° C. Initial RNAi experiments were performed with 10-16 flies/genotype, RNAi-Gal4 and UAS-Dcr controls were run concurrently with each set of RNAi lines. Initial hits were followed up with a second run of at least 10 additional flies.

Circadian rhythms were analyzed in ClockLab Version 6 software (Actimetrics, Wilmette IL). Period, rhythm strength and 24-hour activity were determined for each individual fly using activity data collected from days 2–7 of DD. Period length was determined using ***χ**^2^* periodogram analysis, and relative power (or amplitude) of circadian rhythm was determined using fast Fourier transform (FFT). Each fly’s 24-h activity profile was calculated as the average number of counts per day over 6 days. Fly activity was considered rhythmic if the ***χ**^2^* periodogram showed a peak above the 95% confidence interval and the FFT value was > 0.01 (22). Data from flies that survived the duration of the experiments were pooled and analyzed. Behavioral data were analyzed with one-way analysis of variance (ANOVA). Tukey’s test was used as the post hoc test to compare means between the two control genotypes (flies containing GAL4 or UAS only) and experimental genotype (flies containing both GAL4 and UAS). Differences between groups were considered significant if p < 0.05 by the post hoc test.

#### Behavior experiment: Circadian feeding

Single-fly circadian feeding was assessed using the Automated Recording CAFE assay (43). 5-7 d old male flies were individually housed in plastic chambers containing 300 μL of 2% agar. A 5 μL calibrated capillary containing a liquid diet layered with an infrared-absorbent dye was inserted into each chamber. 2-3 day old male flies were entrained in an LD cycle for at least three days before being transferred to the ARC assay. Flies were then allowed to habituate to the assay for ~18 hours prior to feeding analysis. Capillaries were changed daily throughout the duration of the assay during the morning activity peak.

The liquid diet consisted of 2.5% sucrose, 2.5% yeast extract in sterile water; the infrared-absorbent dye was comprised of 75% mineral oil, 25% dodecane, and 1% Cu (II) 1,4,8,11,15,18,22,25-octabutoxy-29H-phthalocyanine (Sigma-Aldrich, St. Louis, MO). Data was acquired using web cameras (Microsoft LifeCam) modified with an infrared pass filter, through the JavaGrinders-ARC software (http://javagrinders-arc.blogspot.com).

Raw data from JavaGrindersARC acquisition software was processed for noise reduction, identification of feeding bouts and time binning using the Noah python script (https://github.com/HungryFly/flyARC) to generate single fly feeding data in one-minute bins (43). This data was then further processed in Microsoft Excel for averaging of multiple flies of the same genotype. Food consumption per 24-hour day was scored for rhythmicity by JTK_Cycle(53). Statistical analysis of daily consumption was performed in OriginLab 2020.

**Table S1:**
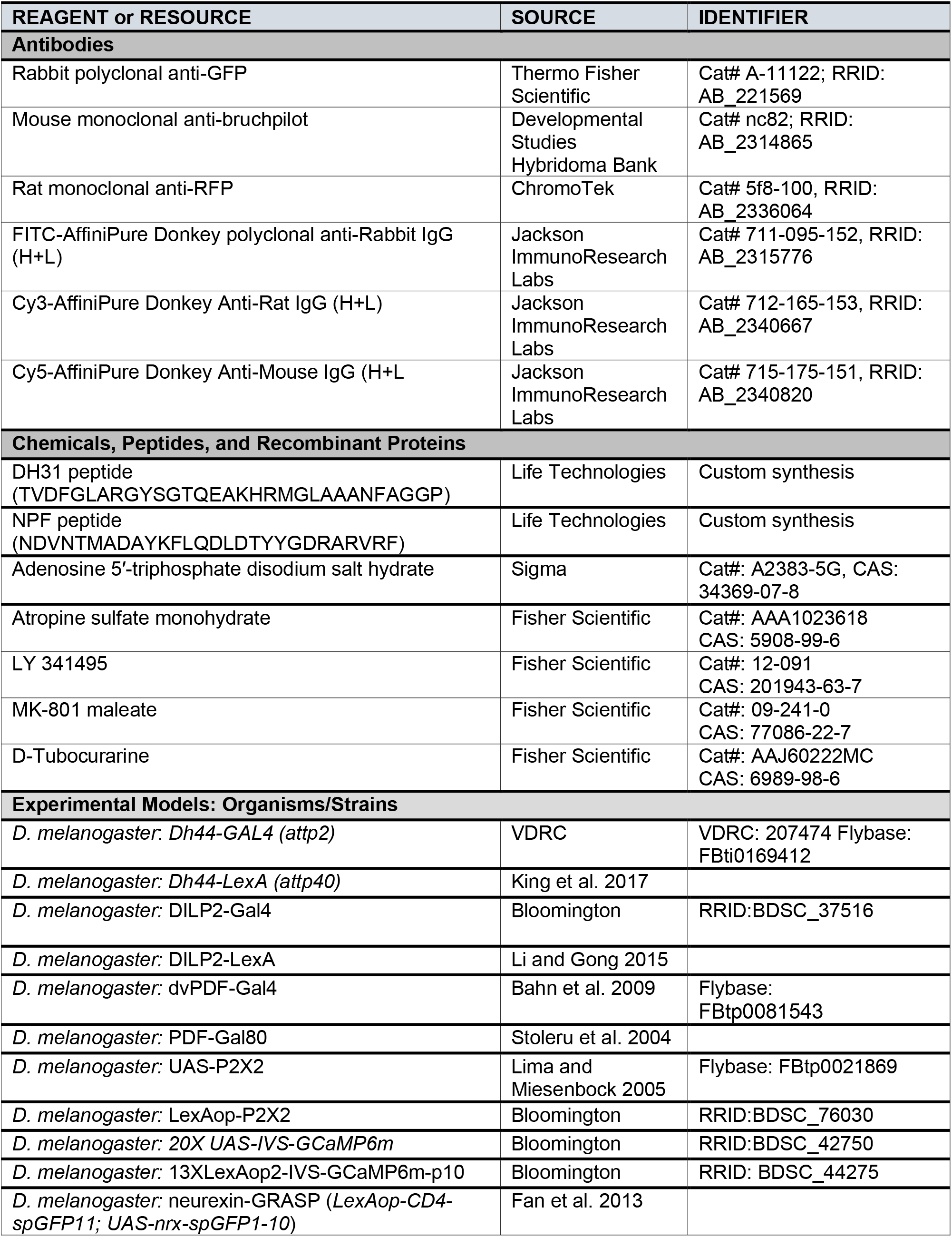
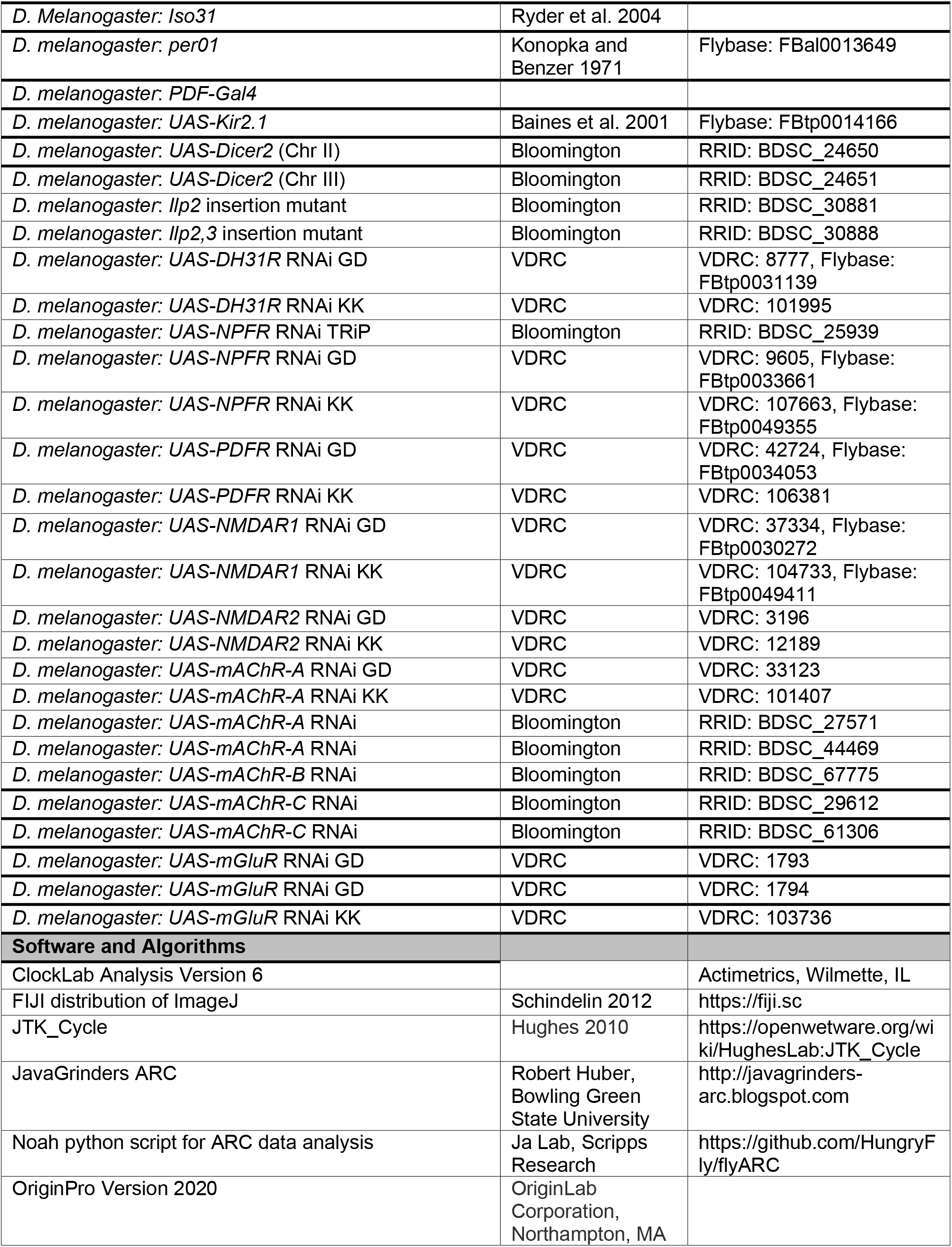
Key Reagents

